# Imputed gene associations identify replicable *trans*-acting genes enriched in transcription pathways and complex traits

**DOI:** 10.1101/471748

**Authors:** Heather E. Wheeler, Sally Ploch, Alvaro N. Barbeira, Rodrigo Bonazzola, Angela Andaleon, Alireza Fotuhi Sishpirani, Ashis Saha, Alexis Battle, Sushmita Roy, Hae Kyung Im

## Abstract

Regulation of gene expression is an important mechanism through which genetic variation can affect complex traits. A substantial portion of gene expression variation can be explained by both local (*cis*) and distal (*trans*) genetic variation. Much progress has been made in uncovering *cis*-acting expression quantitative trait loci (cis-eQTL), but *trans*-eQTL have been more difficult to identify and replicate. Here we take advantage of our ability to predict the *cis* component of gene expression coupled with gene mapping methods such as PrediXcan to identify high confidence candidate *trans*-acting genes and their targets. That is, we correlate the *cis* component of gene expression with observed expression of genes in different chromosomes. Leveraging the shared *cis*-acting regulation across tissues, we combine the evidence of association across all available GTEx tissues and find 2356 *trans*-acting/target gene pairs with high mappability scores. Reassuringly, *trans*-acting genes are enriched in transcription and nucleic acid binding pathways and target genes are enriched in known transcription factor binding sites. Interestingly, *trans*-acting genes are more significantly associated with selected complex traits and diseases than target or background genes, consistent with percolating *trans* effects. Our scripts and summary statistics are publicly available for future studies of *trans*-acting gene regulation.

## Introduction

Transcription is modulated by both proximal genetic variation (cis-acting), which likely affects DNA regulatory elements near the target gene, and distal genetic variation (*trans*-acting). This distal genetic variation likely affects regulation of a transcription factor (or coactivator) that goes on to regulate a target gene, often located on a different chromosome from the transcription factor gene. Expression quantitative trait loci (eQTL) mapping has been successful at identifying and replicating SNPs associated with gene expression in *cis*, typically meaning SNPs within 1 Mb of the target gene. Because effect sizes are large enough, around 100 samples in the early eQTL studies was sufficient to detect replicable associations in the reduced multiple testing space of *cis*-eQTLs (Cheung et al., 2005; Myers et al., 2007; Stranger et al., 2007).

*Trans*-eQTLs have been more difficult to replicate because their effect sizes are usually smaller and the multiple testing burden for testing all SNPs versus all genes can be too large to overcome. A few studies have had some success; one that focused on known GWAS SNPs, with a discovery cohort of 5311 individuals and a replication cohort of 2775 individuals, identified and replicated 103 *trans*-eQTLs in whole blood (Westra et al., 2013). A recent follow-up to this study examined GWAS SNPs in 31,684 individuals and found *trans*-eQTLs in 36% of SNPs tested (Vosa et al., 2018). Unlike *cis*-eQTLs, *trans*-eQTLs are more likely to be tissue-specific, rather than shared across tissues (Aguet et al., 2017; Vosa et al., 2018). However, a large fraction (52%) of *trans*-eQTLs colocalize with at least one *cis*-eQTL signal (Vosa et al., 2018).

Here, we apply PrediXcan (Gamazon et al., 2015) and MultiXcan to map *trans*-acting genes, rather than mapping *trans*-eQTLs (SNPs). Our method provides directionality, that is, whether the *trans*-acting gene activates or represses its target gene. We use genome-transcriptome data sets from the Framingham Heart Study (FHS) (Joehanes et al., 2017), Depression Genes and Networks (DGN) cohort (Battle et al., 2014), and the Genotype-Tissue expression (GTEx) Project (Aguet et al., 2017). We show that our approach, called *trans*-PrediXcan, can identify replicable *trans-acting* regulator/target gene pairs. To leverage sharing of *cis*-eQTLs across tissues and improve our power to detect more *trans*-acting effects, we combine predicted expression across tissues in our *trans*-MultiXcan model and show that it increases significant *trans*-acting/target gene pairs >10-fold.

Pathway analysis reveals the *trans*-acting genes are enriched in transcription and nucleic acid binding pathways and target genes are enriched in known transcription factor binding sites, indicating that our method identifies genes of expected function. We show that *trans*-acting genes are more strongly associated with immune-related traits and height than target or background genes, demonstrating that *trans*-acting genes likely play a key role in the biology of complex traits.

## Methods

### Genome and transcriptome data

#### Framingham Heart Study (FHS)

We obtained genotype and exon expression array data (Joehanes et al., 2017; Zhang et al., 2015) through application to dbGaP accession phs000007.v29.p1. Genotype imputation and gene level quantification were performed by our group previously (Wheeler et al., 2016), leaving 4838 European ancestry individuals with both genotypes and observed gene expression levels for analysis. We used the Affymetrix power tools (APT) suite to perform the preprocessing and normalization steps. First the robust multi-array analysis (RMA) protocol was applied which consists of three steps: background correction, quantile normalization, and summarization (Irizarry et al., 2003). The summarized expression values were then annotated more fully using the annotation databases contained in the huex10stprobeset.db (exon-level annotations) and huex10sttranscriptcluster.db (gene-level annotations) R packages available from Bioconductor. The genotype data were then split by chromosome and pre-phased with SHAPEIT (Delaneau, Marchini, & Zagury, 2012) using the 1000 Genomes phase 3 panel and converted to vcf format. These files were then submitted to the Michigan Imputation Server (https://imputationserver.sph.umich.edu/start.html) (Fuchsberger, Abecasis, & Hinds, 2015; Howie, Fuchsberger, Stephens, Marchini, & Abecasis, 2012) for imputation with the Haplotype Reference Consortium version 1 panel (McCarthy et al., 2016). Approximately 2.5M non-ambiguous strand SNPs with MAF > 0.05, imputation R^2^ > 0.8 and, to match GTEx gene expression prediction models, inclusion in HapMap Phase II were retained for subsequent analyses.

#### Depression Genes and Networks (DGN)

We obtained genotype and whole blood RNA-Seq data through application to the NIMH Repository and Genomics Resource, Study 88 (Battle et al., 2014). For all analyses, we used the HCP (hidden covariates with prior) normalized gene-level expression data used for the trans-eQTL analysis in Battle et al. (Battle et al., 2014) and downloaded from the NIMH repository. Quality control and genotype imputation were performed by our group previously (Wheeler et al., 2016), leaving 922 European ancestry individuals with both imputed genotypes and observed gene expression levels for analysis. Briefly, the 922 individuals were unrelated (all pairwise < 0.05) and thus all included in downstream analyses. Imputation of approximately 650K input SNPs (minor allele frequency [MAF] > 0.05, Hardy-Weinberg Equilibrium [P > 0.05], non-ambiguous strand [no A/T or C/G SNPs]) was performed on the Michigan Imputation Server (Fuchsberger et al., 2015; Howie et al., 2012) with the following parameters: 1000G Phase 1 v3 ShapeIt2 (no singletons) reference panel, SHAPEIT phasing, and EUR population.

Approximately 1.9M non-ambiguous strand SNPs with MAF > 0.05, imputation R^2^ > 0.8 and, to match GTEx gene expression prediction models, inclusion in HapMap Phase II were retained for subsequent analyses.

### Gene expression prediction models

Elastic net (alpha = 0.5) models built using GTEx V6p genome-transcriptome data from 44 tissues (Barbeira et al., 2018) were downloaded from http://predictdb.org/ from the `GTEx-V6p-HapMap-2016-09-08.tar.gz` archive.

### Mappability quality control

Genes with mappability scores less than 0.8 and gene pairs with a positive cross-mappability k-mer count were excluded from our analysis (Saha & Battle, 2018; Saha et al., 2017). Gene mappability is computed as the weighted average of its exon-mappability and untranslated region (UTR)-mappability, weights being proportional to the total length of exonic regions and UTRs, respectively. Mappability of a k-mer is computed as 1/(number of positions k-mer maps in the genome). For exonic regions, k = 75 and for UTRs, k = 36. Cross-mappability between two genes, A and B, is defined as the number of gene A k-mers (75-mers from exons and 36-mers from UTRs) whose alignment start within exonic or untranslated regions of gene B (Saha & Battle, 2018; Saha et al., 2017).

In addition, to further guard against false positives, we retrieved RefSeq Gene Summary descriptions from the UCSC hgFixed database on 2018-10-04 and removed genes from our analyses with a summary that contained one or more of the following strings: “paralog”, “pseudogene”, “retro”.

### *trans*-PrediXcan

In order to map *trans*-acting regulators of gene expression, we implemented *trans*-PrediXcan, which consists of two steps. First, we predict gene expression levels from genotype dosages using models trained in independent cohorts to protect against false positives that may occur by training and testing in the same cohort. As in PrediXcan (Gamazon et al., 2015), this step gives us an estimate of genetic component of gene expression, 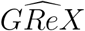, for each gene. In the second step, for each 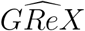 estimate, we calculate the correlation between 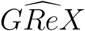 and the observed expression level of each gene located on a different chromosome. As in Matrix eQTL (Shabalin, 2012), variables were standardized to allow fast computation of the correlation and test statistic. In the discovery phase, we predicted gene expression in the FHS cohort using each of 44 tissue models from the GTEx Project. Significance was assessed via the Benjamini-Hochberg false discovery rate (FDR) method (Benjamini & Hochberg, 1995), with FDR < 0.05 in each individual tissue declared significant. We tested discovered *trans*-acting/target gene pairs for replication in the DGN cohort and declared those with P<0.05 replicated. To estimate the expected true positive rate, we calculate *π*_1_ statistics using the qvalue method (Aguet et al., 2017; Storey & Tibshirani, 2003). *π*_1_ is the expected true positive rate and was estimated by selecting the gene pairs with FDR < 0.05 in FHS and examining their P value distribution in DGN. πo is the proportion of false positives estimated by assuming a uniform distribution of null P values and *π*_1_ = 1 — *π_0_* (Storey & Tibshirani, 2003).

For comparison to our *trans*-PrediXcan method, we performed traditional *trans*-eQTL analysis in FHS and DGN using Matrix eQTL (Shabalin, 2012), where *trans* is defined as genes on different chromosomes from each SNP.

### *trans*-MultiXcan

To determine if jointly modeling the genetic component of gene expression across tissues would increase power to detect *trans*-acting regulators, we applied MultiXcan (Barbeira et al., 2019) to our transcriptome cohorts. In our implementation of MultiXcan, predicted expression from all available GTEx tissue models (up to 44) were used as explanatory variables. To avoid multicolinearity, we use the first *k* principal components of the predicted expression in our regression model for association with observed (target) gene expression. We keep the first *k* principal components out of *i* principal components estimated where

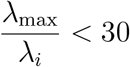

where λ*_i_* is an eigenvalue in the predicted expression covariance matrix (Barbeira et al., 2019). A range of thresholds were previously tested and yielded similar results (Barbeira et al., 2019). We used an F-test to quantify the significance of the joint fit. We tested *trans*-acting/target gene pairs discovered in FHS (FDR < 0.05) for replication in the DGN cohort and declared those with P < 0.05 replicated.

### eQTLGen comparison

We compared our *trans*-PrediXcan and *trans*-MultiXcan FHS results to eQTLs discovered in eQTLGen, a blood eQTL study of 31,684 individuals (Vosa et al., 2018). Note, eQTLGen includes the FHS cohort (n=4838) we used in our *trans*-PrediXcan and *trans*-MultiXcan analyses, and thus it is not a completely independent cohort. To determine the expected distribution of *trans*-eQTLs under the null of no association between predicted and observed expression, we randomly sampled without replacement the lists of predicted and observed genes to generate 1000 sets of “*trans*-acting/target gene pairs”, each the same size and with the same chromosome distribution as the observed results from either *trans*-PrediXcan or *trans*-MultiXcan. We then counted how many “*trans*-acting genes” in each set had an eSNP in their expression prediction model (non-zero effect size) that targeted the same gene in eQTLGen. We compared this distribution to the observed number of *trans*-acting/target gene pairs that had a *trans*-eQTL in eQTLGen to obtain an empirical P value (the number of times the permuted overlap exceeded the observed overlap divided by 1000). To calculate the fold-enrichment of *trans*-eQTLs found in our top *trans*-PrediXcan and *trans*-MultiXcan FHS gene pairs (FDR < 0.05), we determined how many gene pairs included a matching eQTLGen *trans*-eQTL across all tested gene pairs.

### Pathway enrichment analysis

We used FUMA (Functional mapping and annotation of genetic associations) (Watanabe, Taskesen, Van Bochoven, & Posthuma, 2017) to test for enrichment of biological functions in our top *trans*-acting and target genes. We limited our hypergeometric enrichment tests to Reactome (MSigDB v6.1 c2), Gene Ontology (GO) (MMSigDB v6.1 c5), transcription factor targets (MSigDB v6.1 c3), and GWAS Catalog (e91_r2018-02-06) pathways. We required at least 5 *trans*-acting or target genes to overlap with each tested pathway. For the *trans*-acting gene enrichment tests, there were 182 unique *trans*-acting genes at FDR < 0.05 in FHS and P < 0.05 in DGN (Table S2) and the background gene set was the 16,185 genes with a MultiXcan model. For the target gene enrichment tests, there were 211 unique target genes at FDR < 0.05 in FHS and P < 0.05 in DGN (Table S2) and the background gene set was the 12,445 expressed genes. Pathways with Benjamini-Hochberg FDR < 0.05 were considered significant and reported.

We also tested the larger discovery gene sets from FHS (FDR < 0.05) for enrichment in known transcription factors and signaling proteins. The list of transcription factors were collected from Ravasi et al. (Ravasi et al., 2010) and signaling proteins were genes annotated as phosphatases and kinases in Uniprot (Roy et al., 2013; The UniProt Consortium, 2012). We used the hypergeometric test (hypergeom function from scipy.stats Python library) to determine the significance of enrichment. Given the size of the background gene set, *M*, number of genes with the property of interest in the background, *K*, and the size of the selected gene set, *N*, the hypergeometric test calculates the probability of observing *x* or more genes in the selected gene set with the property of interest. In our setting, *K* is the number of genes annotated as a TF or signaling protein and *N* is the size of the discovery gene sets.

### *trans*-acting and target gene association studies with complex traits

We retrieved S-PrediXcan (summary statistic PrediXcan) results from the gene2pheno.org database (Barbeira et al., 2018) for immune-related traits and height. We focused on S-PrediXcan results obtained from gene expression prediction models built using DGN whole blood because that was the largest model cohort with results available. Because the expression prediction models were built using whole blood data, we chose to examine blood and immune related traits available in gene2pheno.org from UK Biobank (UKB) and a second cohort. We also examined height due to the large cohorts available. Traits available from UKB that we analyzed include “50 standing height” (n=500,131), “Non-cancer illness code, self-reported: asthma” (n= 382,462), and “Non-cancer illness code, self-reported: systemic lupus erythematosis/sle” (n= 382,462). Red and white blood cell count S-PrediXcan results were available from a meta-analysis that combined the UKB and INTERVAL cohorts, n=173,480 (Astle et al., 2016). We also examined S-PrediXcan results for systemic lupus erythematosus from IMMUNOBASE (n=23,210) (Bentham et al., 2015), asthma from GABRIEL (n=26,475) (Moffatt et al., 2010), and height from GIANT (n=253,288) (Wood et al., 2014). For each trait, we compared the observed vs. expected P value distributions via QQ plots for three groups of genes: *trans*-acting genes discovered in FHS MultiXcan (FDR < 0.05), target genes discovered in FHS MultiXcan (FDR < 0.05), and background genes tested in MultiXcan that were not significant. In each cohort, there were approximately 560 *trans*-acting genes (FHS FDR < 0.05), 700 target genes (FHS FDR < 0.05), and 9900 background genes.

### R packages

R packages used in this work include huex10stprobeset.db (MacDonald, 2015a), huex10sttranscriptcluster.db (MacDonald, 2015b), MatrixEQTL (Shabalin, 2012), qvalue (Bass, Storey, Dabney, & Robinson, 2017; Storey & Tibshirani, 2003), data.table (Dowle & Srinivasan, 2017), dplyr (Wickham, Francois, Henry, & Muller, 2017), ggplot2 (Wickham, 2009), ggrepel (Slowikowski, 2017), readxl (Wickham & Bryan, 2017), and gridExtra (Auguie, 2017).

## Results

### *Trans*-acting gene discovery and validation with *trans*-PrediXcan

We sought to map *trans*-acting and target gene pairs by applying the PrediXcan framework to observed expression as traits and term the approach *trans*-PrediXcan (Fig. 1). We excluded genes with poor genome mappability from our analyses (see Methods). We compared *trans*-PrediXcan results between the discovery FHS whole blood cohort (n = 4838) and the validation DGN whole blood cohort (n = 922). We first used PrediXcan (Gamazon et al., 2015) to generate a matrix of predicted gene expression from FHS genotypes using prediction models built in GTEx whole blood (Barbeira et al., 2018). Then, we calculated the correlation between predicted and observed FHS whole blood gene expression. Examining the correlations of gene pairs on different chromosomes, 55 pairs were significantly correlated in FHS, with an expected true positive rate (*π*_1_) of 0.72 in DGN (Table 1, Fig. S1). Gene pair information and summary statistics are shown in Table S1.

**Figure 1.**
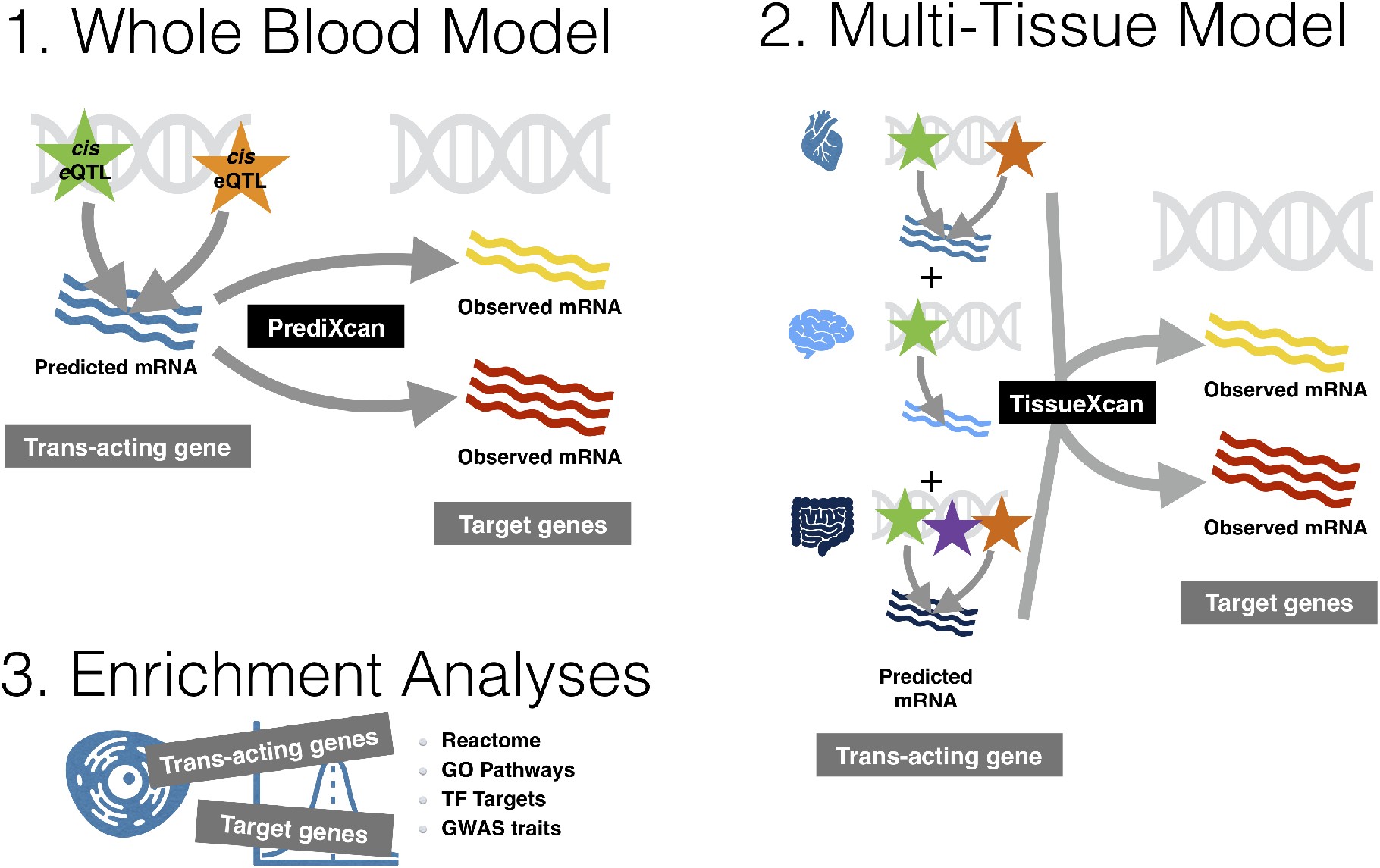
Overview of approach to detect and characterize *trans*-acting genes. First, in our Whole Blood Model, we predict mRNA expression levels from *cis* region eQTLs, using weights trained in a single tissue (GTEx whole blood). These predicted expression levels *(trans*-acting genes) are tested for association with observed expression on different chromosomes (target genes). Second, in our Multi-Tissue Model, we use predicted mRNA expression levels from multiple tissues in a multiple regression to detect *trans*-acting genes and their targets. Third, we compare models and test significant *trans*-acting and target genes for enrichment in pathways or in GWAS traits.

**Table 1.** *Trans*-acting and target gene pair counts and replication rates across GTEx tissue models.

| Model | FHS FDR<0.05 | FHS Tested | DGN P<0.05 | DGN Tested | DGN $\pi_1$ |
| --- | --- | --- | --- | --- | --- |
| Multi-Tissue<br>(MultiXcan) | 2356 | 2.0E+08 | 535 | 1902 | 0.49 |
| Whole Blood<br>(PrediXcan) | 55 | 2.4E+07 | 26 | 54 | 0.72 |

FHS = Framingham Heart Study whole blood cohort, DGN = Depression Genes and Networks whole blood cohort, P = p-value, FDR = Benjamini-Hochberg false discovery rate, $\pi_1$ = expected true positive rate.

Of the 55 *trans*-acting/target gene pairs, 29 had a negative effect size, meaning the *trans*-acting gene may be a repressor because decreased expression of the *trans*-acting gene is associated with increased expression of the target gene. Conversely, 26 had a positive effect size, meaning the *trans*-acting gene may be an activator because increased expression of the *trans*-acting gene is associated with increased expression of the target gene. Note that the directions of effect of 69% of these gene pairs discovered in FHS are consistent in DGN (Fig. 2). None of the *trans*-acting/target gene pairs we identified also acted in the reverse direction, that is, if gene A was *trans*-acting to target gene B, gene B was not also *trans*-acting to target gene A. Looking at all results, beyond just the top signals, there was no correlation in effect sizes between such pairs (P = 0.53). Therefore, our *trans*-PrediXcan method is not simply capturing a co-expression network.

**Figure 2.**
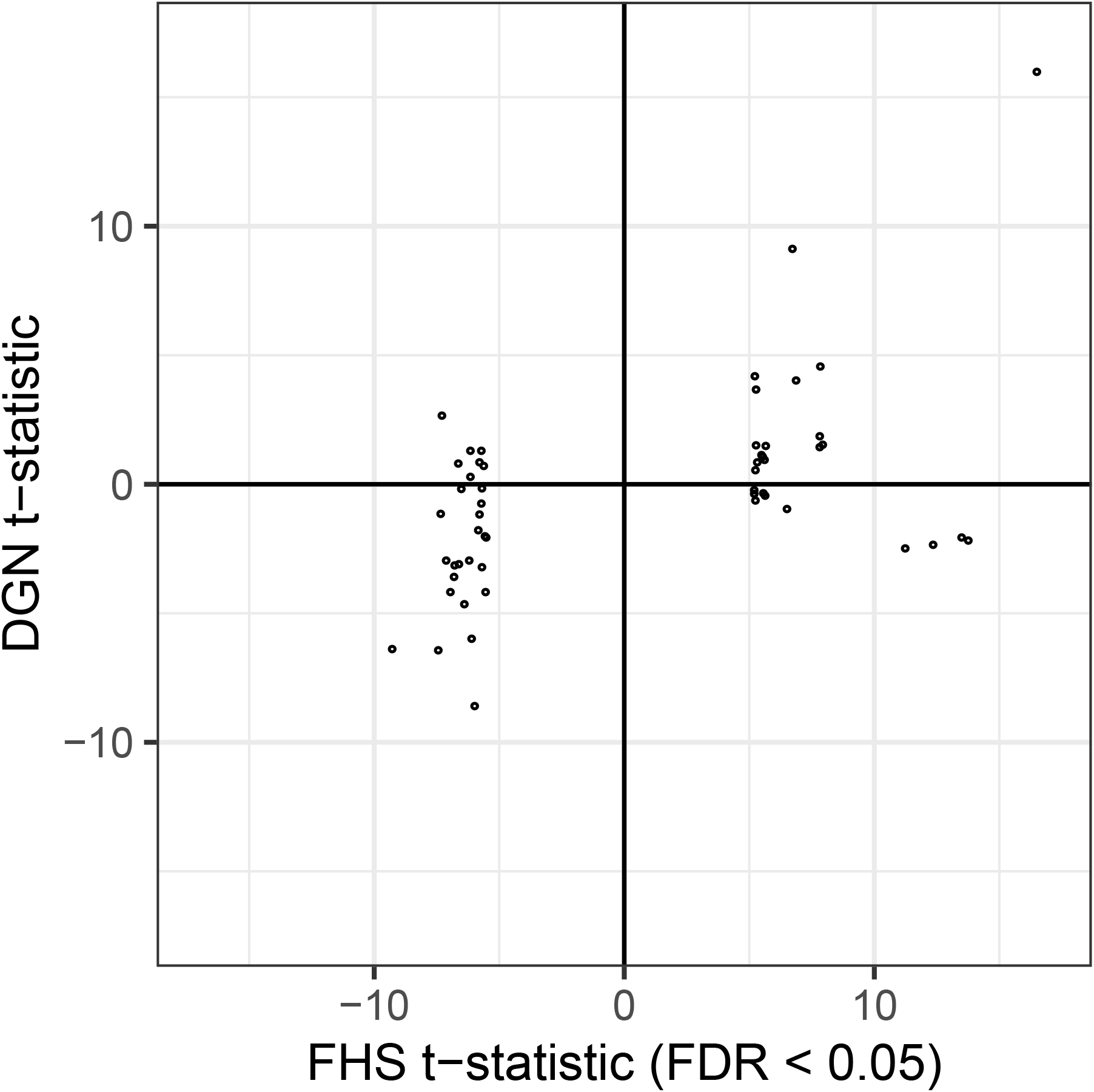
Comparison between FHS and DGN results using the GTEx whole blood prediction models. Results of *trans*-acting gene pairs with FDR < 0.05 in the discovery cohort (FHS) are shown for both FHS (x-axis) and the validation cohort DGN (y-axis). The t-statistics from the linear models testing predicted *trans*-acting expression for association with observed target gene expression are plotted.

To compare the performance of our *trans*-PrediXcan approach to traditional *trans*-eQTL analysis, we also examined the P-value distribution of top FHS *trans*-eQTLs (FDR < 0.05) in DGN to determine the expected true positive rate. In our SNP-level *trans*-eQTL analysis, *π*_1_ was 0.46, 36% lower than the *trans*-PrediXcan *π*_1_ of 0.72. We also compared our results to a recent blood eQTL study in the eQTLGen cohort (Vosa et al., 2018). Of the 55 whole blood model gene pairs we discovered in FHS, 5/55 (9%) have at least one *trans*-eQTL (FDR < 0.05) shared with eQTLGen, more than expected by chance based on the genes tested (empirical P < 0.001, Table S1). This means our prediction model for the *trans*-acting gene includes a nonzero weight for the eQTLGen eSNP and that the target gene in eQTLGen and our whole blood results is the same. Across all 2.4×10^7^ gene pairs tested, just 3547 (0.01%) included a shared *trans*-eQTL with eQTLGen. Thus, top *trans*-PrediXcan gene pairs show a 900-fold enrichment (9/0.01) of eQTLGen *trans*-eQTLs among whole blood model prediction SNPs compared to all gene pairs tested. In addition, of the 5 gene pairs with a matching *trans*-eQTL in eQTLGen, all 5 also had a *cis*-eSNP in eQTLGen (FDR < 0.05) targeting the *trans*-acting gene from our results and present in the prediction model of the *trans*-acting gene. A list of these overlapping eSNPs is shown in Table S2.

### Multi-tissue prediction improves *trans*-acting gene discovery and validation

To leverage tissue sharing of *cis*-eQTLs, we used a multivariable regression approach called MultiXcan, which accounts for correlation among predicted expression levels across 44 GTEx tissues (Barbeira et al., 2019). Notice that even though we seek to detect *trans* regulation, the instruments we are using, i.e. predicted expression, are based on *cis* regulation. Thus, it makes sense to combine information across tissues to obtain the best local predictor of gene expression. To address multicolinearity issues, MultiXcan uses principal component analysis to reduce the number of independent variables to those with the largest variation (Barbeira et al., 2019). When we applied *trans*-MultiXcan to the FHS data, the number of *trans*-acting/target gene pairs increased dramatically (Fig. 3). At FDR < 0.05, there were 2,356 *trans*-acting gene pairs discovered in FHS using the multi-tissue method, while only 55 pairs were discovered with the GTEx whole blood predictors alone (Table 1). We could test 1,902 of these multi-tissue gene pairs for replication in DGN and found 535 of them were significant at P < 0.05 (blue in Fig. 4). Although the expected true positive rate was lower with the MultiXcan model (π_1_ = 0.49) than with the single tissue model (π_1_ = 0.72), the absolute number of replicate gene pairs was much higher (Table 1, Fig. S1). Thus, the number of genes that replicated in both cohorts was 20 times higher in the multi-tissue model compared to the whole blood model (Table 1). Similarly, for gene pairs tested in both models, the adjusted R^2^ was consistently higher in the multi-tissue model than the whole blood model across gene pairs (Fig. S2). Summary statistics of the 2,356 gene pairs discovered in the *trans*-MultiXcan are available in Table S3.

**Figure 3.**
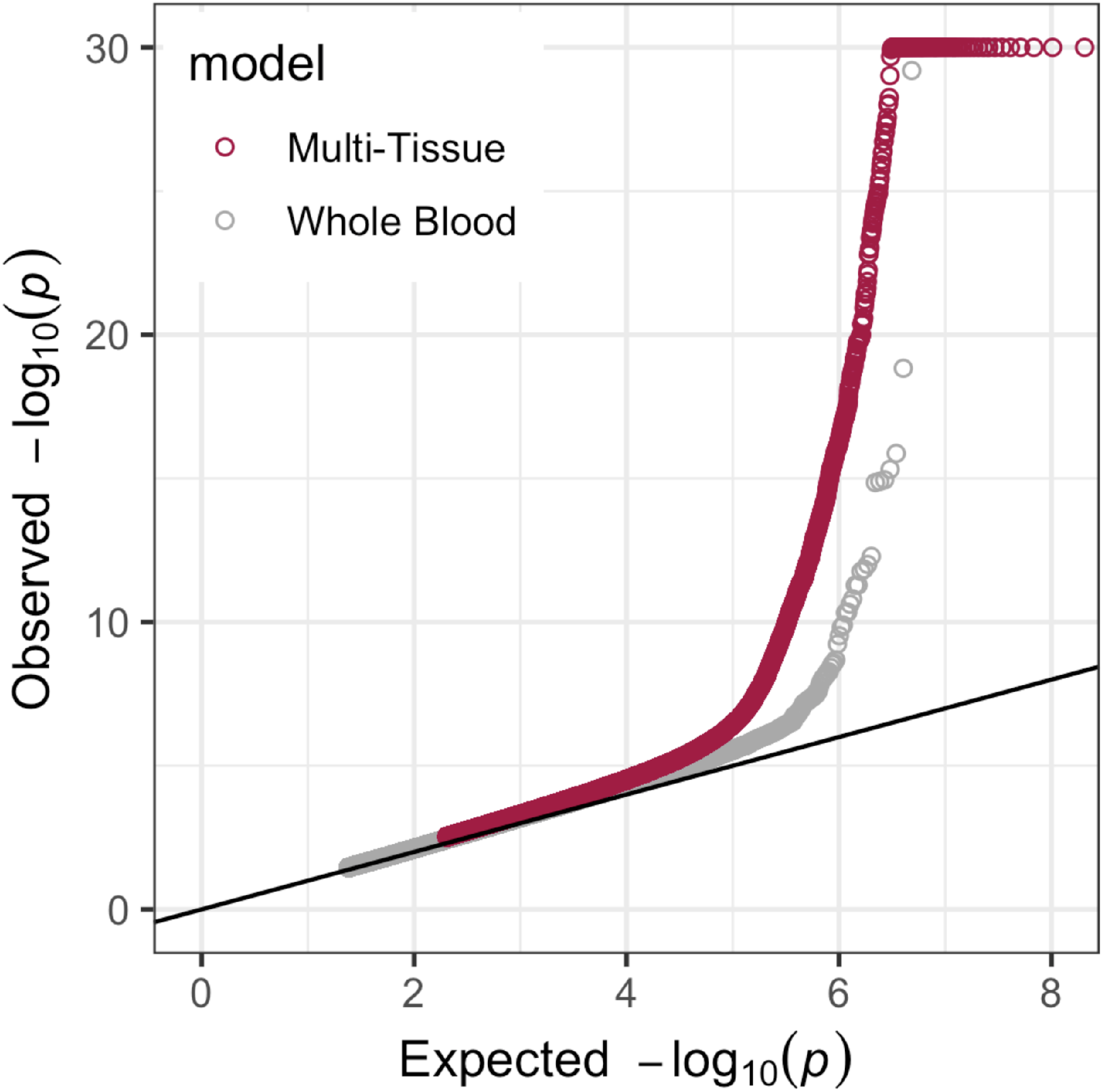
Multi-Tissue *trans*-MultiXcan finds more *trans*-acting gene pairs than a single tissue *trans*-PrediXcan (Whole Blood) model. Quantile-quantile plots show an increase in signal in the Multi-Tissue model compared to the Whole Blood model. -log_10_ p-values are capped at 30 for ease of viewing. The 1e6 most significant P values in each model are plotted to manage file size.

**Figure 4.**
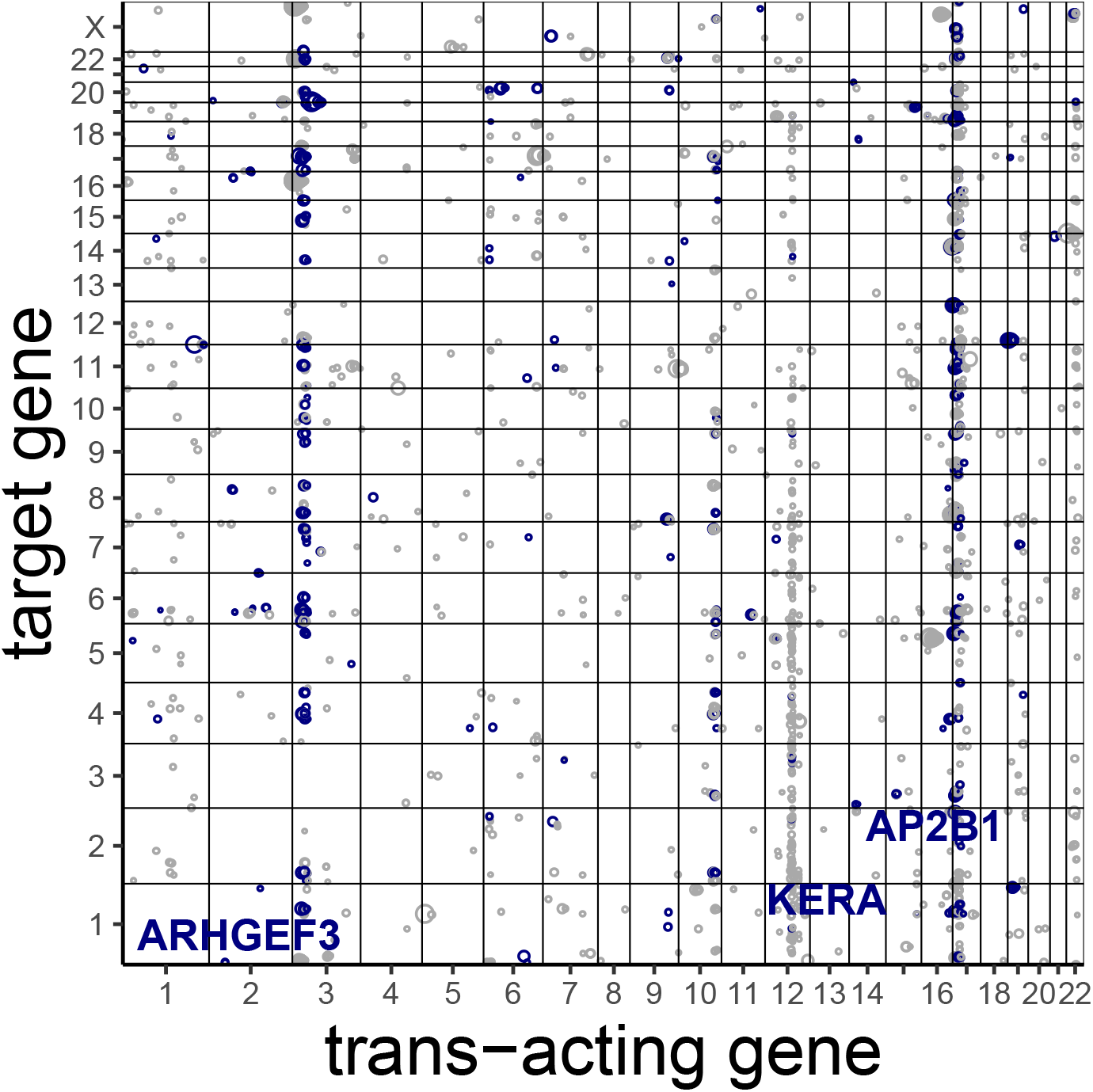
*trans*-acting/target gene pairs discovered using MultiXcan in FHS. Each point corresponds to one gene pair (FHS FDR < 0.05) positioned by chromosomal location of the *trans*-acting gene (x-axis) and target gene (y-axis). Size of the point is proportional to the -logı_0_ p-value in FHS. Gene pairs that replicated in DGN MultiXcan (P < 0.05) are colored blue. Master *trans*-acting loci with greater than 50 target genes are labeled.

Of the MultiXcan gene pairs we found, 728/2356 (31%) replicated in the blood eQTLGen cohort. That is, 31% of MultiXcan gene pairs have at least one *trans*-eQTL shared with eQTLGen, more than expected by chance based on the genes tested (empirical P < 0.001, Table S3). This means that at least one tissue’s prediction model for the *trans*-acting gene includes a nonzero weight for the eQTLGen eSNP and that the target gene in eQTLGen and our multi-tissue results is the same. Across all 2×10^8^ gene pairs tested, 168,893 (0.08%) included a shared *trans*-eQTL with eQTLGen. Thus, top *trans*-MultiXcan gene pairs show an approximately 400-fold enrichment (31/0.08) of eQTLGen *trans*-eQTLs among prediction SNPs compared to all gene pairs tested. *trans*-eQTLs with eSNPs in our MultiXcan *trans*-acting gene models with the same target genes are shown in Table S4. In addition, of these 728 gene pairs with a matching *trans*-eQTL in eQTLGen, 283 (39%) also had a *cis*-eSNP in eQTLGen (FDR < 0.05) targeting the *trans*-acting gene from our results and present in the prediction model of the *trans*-acting gene in at least one tissue.

### Master *trans*-acting genes associate with many targets

Points that form vertical lines in Figure 4 are indicative of potential master regulators, i.e. genes that regulate many downstream target genes. We defined master regulators as *trans*-acting genes that associate with 50 or more target genes. In our MultiXcan analysis, we discovered three potential master regulator loci, which are labeled in Figure 4. The most likely master regulator we identified with MultiXcan is *ARHGEF3* on chromosome 3. *ARHGEF3* associated with 53 target genes in FHS (FDR < 0.05) and 45/51 tested replicated in DGN (P < 0.05). Also, SNPs in *ARHGEF3* have previously been identified as *trans*-eQTLs with multiple target genes. *ARHGEF3* encodes a ubiquitously expressed guanine nucleotide exchange factor. Multiple GWAS and functional studies in model organisms have implicated the gene in platelet formation (Astle et al., 2016; Gieger et al., 2011; Schramm et al., 2014; Yao et al., 2017; Zhang et al., 2014). Similarly, SNPs at the chromosome 17 locus we identified have also been identified as *trans*-eQTLs (Kirsten et al., 2015) and one study showed the the *trans* effects are mediated by *cis* effects on *AP2B1* expression (Yao et al., 2017). *AP2B1* encodes a subunit of the adaptor protein complex 2 and GWAS have implicated it in red blood cell and platelet traits (Astle et al., 2016).

### *trans*-acting genes are enriched in transcription factor pathways

We tested replicated *trans*-acting genes for enrichment in Reactome (MSigDB v6.1 c2), Gene Ontology (GO, MSigDB v6.1 c5), transcription factor targets (MSigDB v6.1 c3), and GWAS Catalog (e91_r2018-02-06) pathways using FUMA (Watanabe et al., 2017). In our MultiXcan analysis, there were 174 unique *trans*-acting genes at FDR < 0.05 in FHS and P < 0.05 in DGN (Table S2). We required at least 5 *trans*-acting genes to overlap with each tested pathway. The background gene set used in the enrichment test were the 15,432 genes with a MultiXcan model. All pathways with FDR < 0.05 are shown in Table 2 and their gene overlap lists are available in Table S5.

**Table 2.** Replicated *trans*-acting genes (MultiXcan FDR < 0.05 in FHS and P < 0.05 in DGN) are enriched in transcription and GWAS pathways.

| Source | GeneSet | N | n | P-value | adjusted P |
| --- | --- | --- | --- | --- | --- |
| GO molecular functions | GO nucleic acid binding transcription factor activity | 1000 | 33 | 6.40e-09 | 5.76e-06 |
| Reactome | Reactome generic transcription pathway | 292 | 15 | 2.18e-07 | 1.47e-04 |
| GWAS catalog | Reticulocyte count | 138 | 10 | 4.80e-07 | 1.47e-04 |
| GWAS catalog | Reticulocyte fraction of red cells | 147 | 9 | 6.76e-06 | 2.75e-03 |
| GWAS catalog | White blood cell count | 152 | 8 | 5.90e-05 | 1.57e-02 |
| GWAS catalog | Neuroticism | 134 | 7 | 1.44e-04 | 2.20e-02 |
| GWAS catalog | Platelet count | 228 | 9 | 2.78e-04 | 3.40e-02 |
| GWAS catalog | Crohn's disease | 525 | 15 | 3.19e-04 | 3.54e-02 |

FHS = Framingham Heart Study, DGN = Depression Genes and Networks cohort, N = number of genes in GeneSet tested for *trans*-acting effects, n = number of replicated genes in GeneSet, P-value = FUMA (Watanabe et al., 2017) enrichment P, adjusted P = enrichment Benjamini-Hochberg false discovery rate

The top two most significant pathways were the GO nucleic acid binding transcription factor activity pathway and the reactome generic transcription pathway (Table 2). The *trans*-acting genes in each pathway are spread across multiple chromosomes as shown in Figure S3. *PLAGL1*, which encodes a C2H2 zinc finger protein that functions as a suppressor of cell growth, is a notable *trans*-acting gene in the GO nucleic acid binding transcription factor activity pathway. Of the four *PLAGL1* target genes discovered in FHS, three replicated in DGN (Table S2). One notable gene in the reactome generic transcription pathway is *MED24.* In our MultiXcan analysis, *MED24* targeted 13 genes in FHS (FDR < 0.05) and 8/12 replicated in DGN (P < 0.05, Table S2). *MED24* encodes mediator complex subunit 24. The mediator complex is a transcriptional coactivator complex required for the expression of almost all genes. The mediator complex is recruited by transcriptional activators or nuclear receptors to induce gene expression, possibly by interacting with RNA polymerase II and promoting the formation of a transcriptional pre-initiation complex (Gustafsson & Samuelsson, 2001).

We also found a significant enrichment of transcription factors from Ravasi et al. (Ravasi et al., 2010) in the 766 unique *trans*-acting genes discovered in FHS with FDR < 0.05 (hypergeometric test P = 9.56×10^−3^). However, the same *trans*-acting genes were not enriched in signaling proteins (P = 0.71).

### Target genes are enriched in transcription factor binding sites

We tested MultiXcan replicated target genes for enrichment in the same pathways tested in the *trans*-acting gene analysis. There were 201 unique target genes at FDR < 0.05 in FHS and P < 0.05 in DGN (Table S2). While just eight pathways were enriched in *trans*-acting genes, 118 pathways were enriched in the target genes (Table S5). Two of these 118 target gene enriched pathways were transcription factor binding sites (Table 3). No binding motifs were enriched in the *trans*-acting genes. Additional pathways enriched in target genes included several platelet activation and immune response pathways (Table S5). Target genes were spread across multiple chromosomes (Fig. S4). The target genes were not enriched for reactome generic transcription or GO nucleic acid binding transcription factor activity pathways. The 945 unique target genes discovered in FHS with FDR <0.05 were also not enriched for transcription factors (hypergeometric test P = 0.98) or signaling proteins (P = 0.46) from Ravasi et al. (Ravasi et al., 2010).

**Table 3.** Replicated target genes (MultiXcan FDR < 0.05 in FHS and P < 0.05 in DGN) are enriched in transcription factor (TF) binding sites in the regions spanning up to 4 kb around their transcription starting sites (MSigDB v6.1 c3).

| TF binding site | N | n | P-value | adjusted P |
| --- | --- | --- | --- | --- |
| WGGAATGY_TEF1_Q6 | 247 | 14 | 2.23e-05 | 1.37e-02 |
| PAX8_B | 68 | 6 | 1.56e-04 | 4.79e-02 |

FHS = Framingham Heart Study, DGN = Depression Genes and Networks cohort, N = number of genes in GeneSet tested for target gene effects, n = number of replicated genes in GeneSet, P-value = FUMA (Watanabe et al., 2017) enrichment P, adjusted P = enrichment Benjamini-Hochberg false discovery rate

### *Trans*-acting genes are more likely to associate with complex traits

*Trans*-acting genes may drive complex trait inheritance, which has been formalized in the omnigenic model (Boyle et al., 2017; Liu, Li, & Pritchard, 2018). If true, we hypothesized that the *trans*-acting genes we discovered using our *trans*-MultiXcan model should be more significantly associated with complex traits than both their targets and other background genes. We focused on immune related complex traits because our observed gene expression data in FHS and DGN are from whole blood. We also used height as a representative complex trait because of the large sample sizes available.

Using height and immune related phenotypes from the UK Biobank and other large consortia (see Methods) as representative complex traits, we compared PrediXcan results among three classes of gene: *trans*-acting, target, and background genes. *trans*-acting and target genes were those discovered in our FHS MultiXcan analysis (FDR < 0.05). Background genes are those tested in MultiXcan, but not found significant. We examined QQ plots of PrediXcan results for each class in two large studies of height, red and white blood cell counts, two studies of systemic lupus erythematosus, and two studies of asthma. For each trait, we found that *trans*-acting gene associations are more significant than background gene associations (Fig. 5). Though attenuated in comparison to *trans*-acting genes, target genes are also more significant than background genes for several traits (Fig. 5).

**Figure 5.**
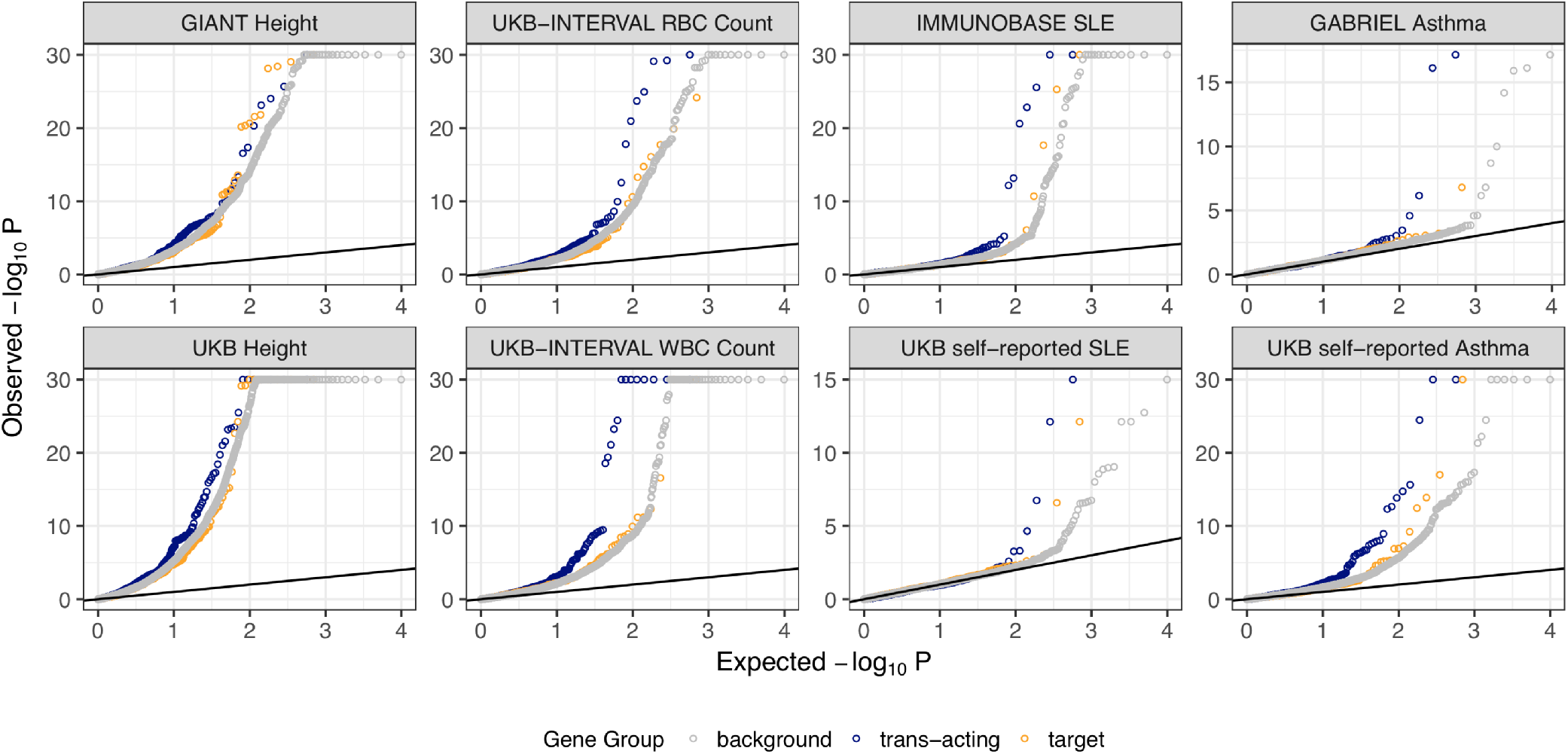
Complex trait associated genes are enriched for *trans*-acting genes. Quantile-quantile plots of S-PrediXcan results for each labeled trait show an increase in signal for *trans*-acting genes (FHS MultiXcan FDR < 0.05) compared to target genes (FHS MultiXcan FDR < 0.05) and background (tested in MultiXcan, but not significant) genes. When present, -log_10_ p-values greater than 30 are capped at 30 for ease of viewing. RBC = red blood cell, WBC = white blood cell, SLE = systemic lupus erythematosus.

## Discussion

We apply the PrediXcan framework to gene expression as a trait (*trans*-PrediXcan approach) to identify *trans*-acting genes that potentially regulate target genes on other chromosomes. We identify replicable predicted gene expression and observed gene expression correlations between genes on different chromosomes. Compared to *trans*-eQTL studies performed in the same cohorts, our *trans*-PrediXcan model shows a higher replication rate for discovered associations. For example, using the GTEx whole blood prediction model we show the expected true positive rate is 0.72 (Table 1). When we performed a traditional *trans*-eQTL study and examined the P-value distribution of top FHS eQTLs (FDR < 0.05) in DGN, the true positive rate was only 0.46. In an independent analysis of the same data, only 4% of eQTLs discovered in FHS replicated in DGN (Joehanes et al., 2017).

In contrast to our results, a recent study concluded *trans*-eQTLs have limited influence on complex trait biology (Yap et al., 2018). However, the authors mention limited power in their analyses and found most of the *trans*-eQTLs examined were not also *cis*-eQTLs for nearby genes (Yap et al., 2018). To combat lack of power, others have used *cis*-mediation analysis to identify *trans*-eQTLs (Yang et al., 2017; Yao et al., 2017). Similar to our approach, a mechanism is built in to significant associations found via *cis*-mediation studies: the *cis*-acting locus causes variable expression of the local gene, which in turn leads to variable expression of its target gene on a different chromosome. Unlike *cis*-mediation analysis, our *trans*-PrediXcan approach allows multiple SNPs to work together to affect expression of the *trans*-acting gene and thus may reveal additional associations. A similar method, developed in parallel to ours, combines *cis*-region SNPs using a cross-validation BLUP to identify *trans*-acting genes within one eQTL cohort (Liu, Mefford, et al., 2018). Our findings have the advantage of discovery in a larger cohort, multiple tissue integration, and replication in an independent cohort.

When predictive models built in 44 different tissues are combined with MultiXcan, we increase the number of *trans*-acting gene pairs identified in FHS and replicated in DGN 20-fold compared to single-tissue models (Table 1). In the recent release of eQTLGen, the largest *trans*-eQTL study to date, 52% of *trans*-eQTL signals colocalize with at least one *cis*-eQTL signal (Vosa et al., 2018). As currently implemented, our *trans*-PrediXcan method will only find gene pairs that have *cis*-acting regulation of the predicted (*trans*-acting) gene. The SNPs used to predict expression of each gene are all within 1Mb of the gene, i.e. in *cis.* Previous work has shown that *cis*-eQTLs are often shared across many tissues (Aguet et al., 2017). Thus, we show combining *cis*-acting effects across tissues as “replicate experiments” increases our power to detect *trans*-acting associations. For example, if there is a *cis*-acting effect that is common across most tissues but the *trans*-acting effect occurs in one specific tissue, MultiXcan will be able to identify the *trans*-acting effect even if we do not have a prediction model in the causal tissue. Our choice to use PC regression is a conservative approach, discarding less informative components of expression variation at the cost of slightly reduced power. This “denoising” property may limit our ability to detect tissue-specific effects, which may be revealed in future studies with larger sample sizes and prediction modeling approaches that include distal genetic variation. Another limitation is that our approach can detect false positives due to linkage disequilibrium and thus colocalization and functional studies are required to reveal the causal *trans*-acting regulator of gene expression.

We found *trans*-acting genes discovered in our MultiXcan analysis were enriched in transcription pathways and thus previously known to function in transcription regulation. Master regulators revealed by MultiXcan, *ARHGEF3* and *AP2B1*, were also previously known (Kirsten et al., 2015; Yao et al., 2017). Our transcriptome association scan presented here integrates gene expression prediction models from multiple tissues and replicates results in an independent cohort. Encouragingly, the *trans*-acting and target genes we identify are enriched in transcription and transcription factor pathways.

Using asthma, lupus, blood cell counts, and height as representative complex traits, *trans*-acting gene associations with these traits are more significant than target and background gene associations in multiple cohorts. This suggests percolating effects of *trans*-acting genes through target genes. We make our scripts and summary statistics available for future studies of *trans*-acting gene regulation at https://github.com/WheelerLab/trans-PrediXcan.

---

**S1 Figure.**
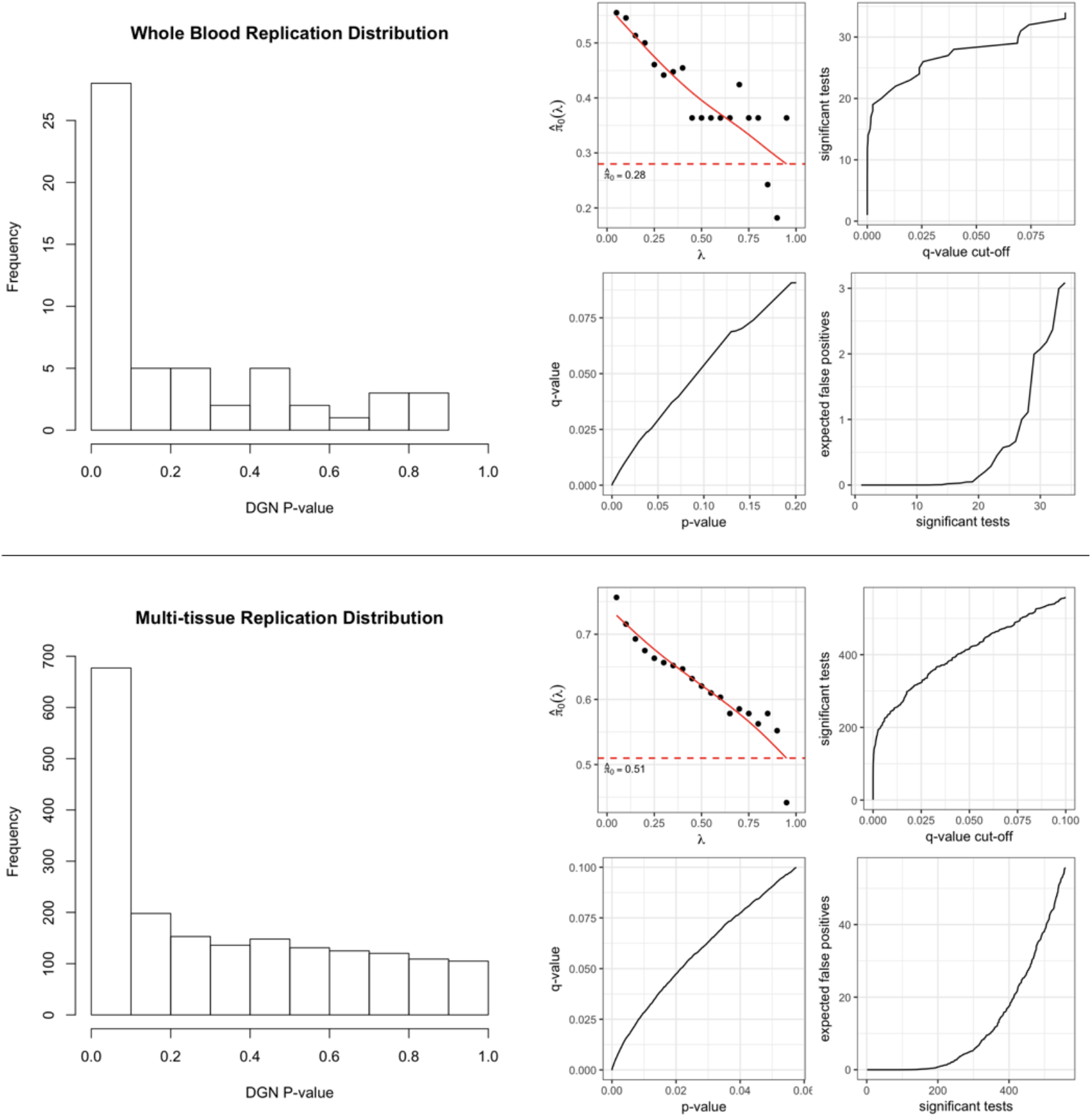
*trans*-acting/target gene pairs meeting significance in FHS (FDR < 0.05) were tested for replication in DGN. Shown are the p-value distributions of these gene pairs in DGN along with *π*_1_ diagnosis plots for the whole blood and multi-tissue models.

**S2 Figure.**
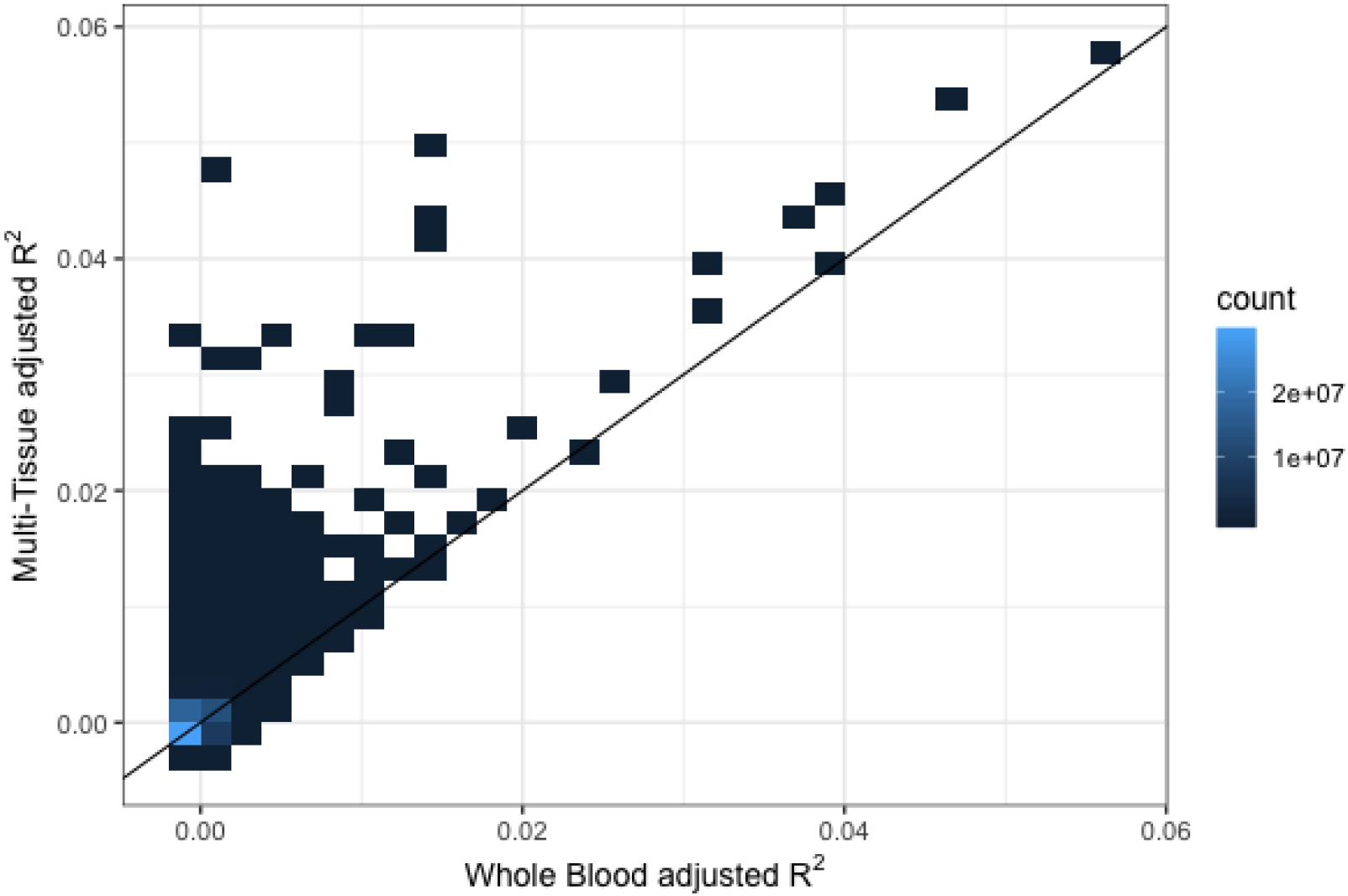
The percent variance explained by *trans*-acting genes is higher in the multi-tissue model than the single tissue whole blood model. Adjusted *R*^2^ values from the multi-tissue model (y-axis) are compared to the adjusted *R*^2^ values from the single tissue model (x-axis) for *trans*-acting/target gene pairs tested in both models.

**S3 Figure.**
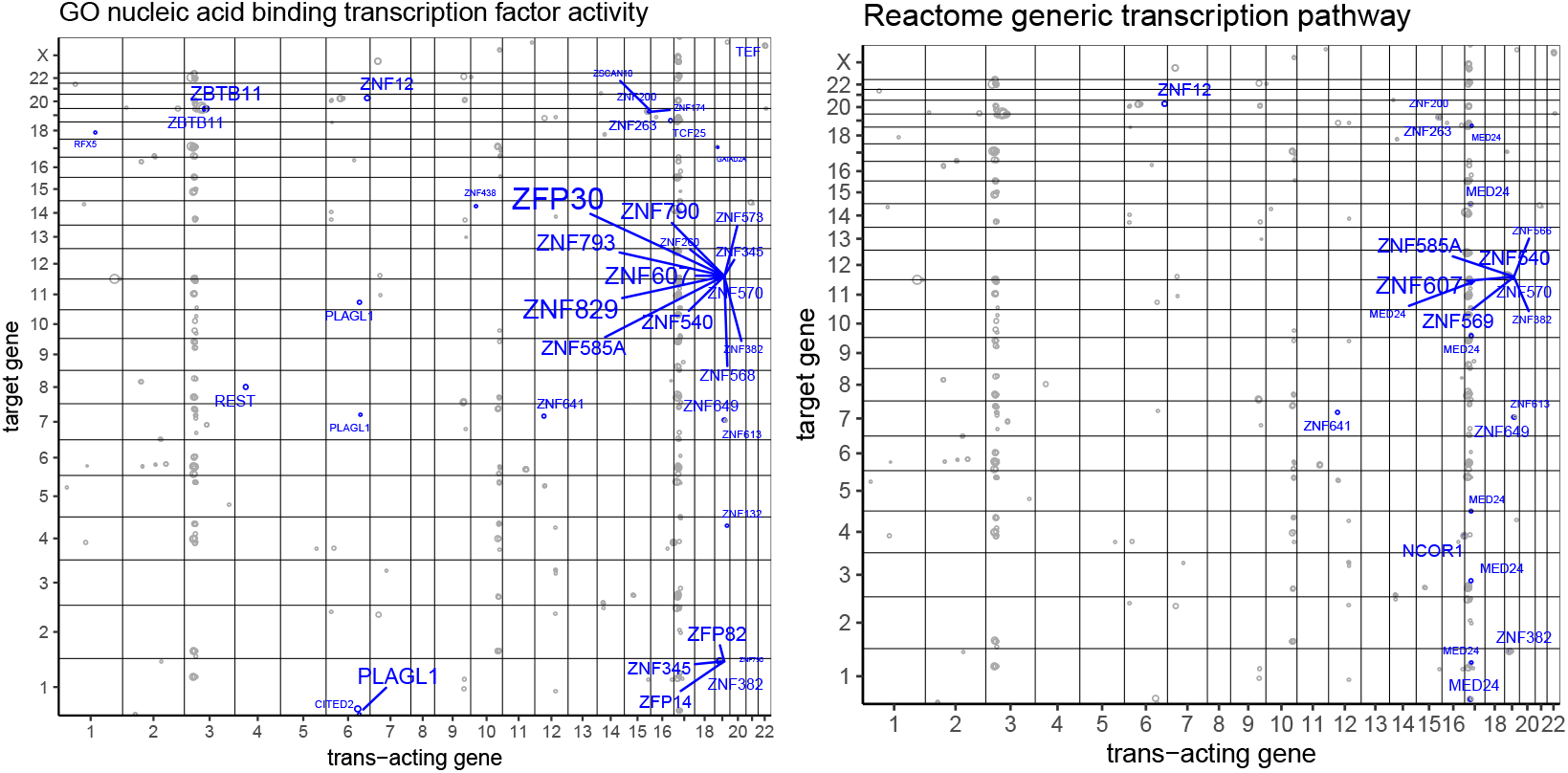
*trans*-acting genes discovered and replicated with MultiXcan are enriched in transcription pathways. Shown are the MultiXcan replicated (FHS FDR <0.05 and DGN P <0.05) results plotted by chromosomal position. The size of the point is proportional to -log_ıν_ p-value. **trans*-acting* genes in the title pathway are labeled in blue.

**S4 Figure.**
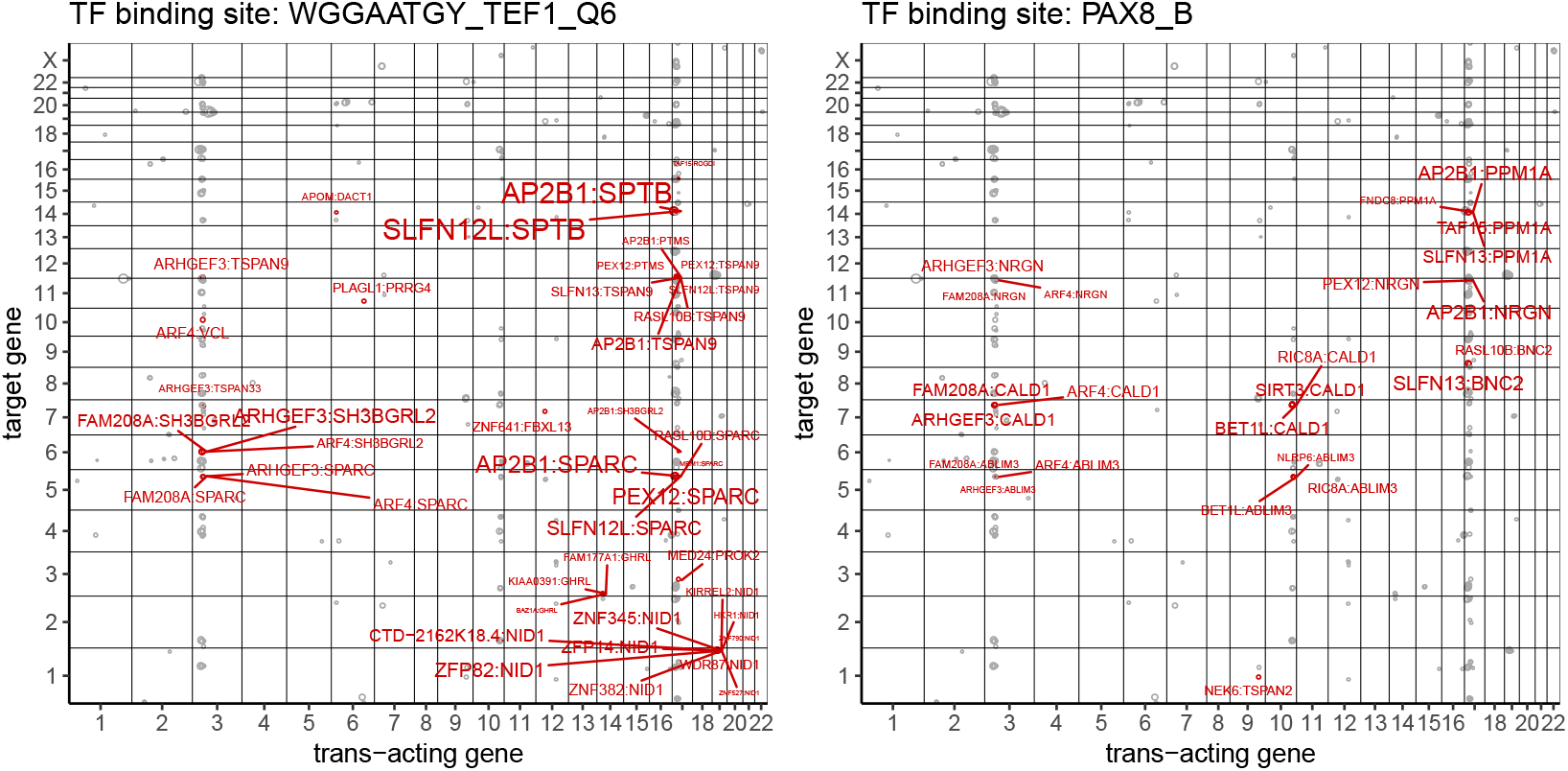
Target genes discovered and replicated with MultiXcan are enriched in transcription factor binding site pathways. Shown are the MultiXcan replicated (FHS FDR <0.05 and DGN P <0.05) results plotted by chromosomal position. The size of the point is proportional to -log_10_ p-value. Gene pairs with the target gene containing the title binding site are labeled in red (trans-acting gene:target gene).

**S1 Table** *trans*-PrediXcan whole blood model results in FHS (FDR < 0.05) and DGN.

**S2 Table** eQTLGen (Vosa et al., 2018) replication of FHS *trans*-PrediXcan whole blood model gene pairs (FDR < 0.05).

**S3 Table** *trans*-MultiXcan results in FHS (FDR < 0.05) and DGN.

**S4 Table** eQTLGen (Vosa et al., 2018) replication of FHS *trans*-MultiXcan whole blood model gene pairs (FDR < 0.05).

**S5 Table** FUMA gene set enrichment results of replicated (FHS FDR < 0.05 and DGN P < 0.05) *trans*-acting and target genes.

